# Enhancing Missense Variant Classification in Predicted Intrinsically Disordered Regions

**DOI:** 10.1101/2025.08.08.669269

**Authors:** Rohan D Gnanaolivu, Steven N Hart

## Abstract

The accurate classification of missense variants is a fundamental challenge in genomics, particularly for those within intrinsically disordered regions (IDRs) where the performance of existing computational predictors is suboptimal. To address this, we developed a machine learning model that extends traditional missense tools with properties that infer globular IDR conformation, phase separation, and protein embeddings. Using ClinVar variant classifications as ground truth, AlphaMissense, EVE, and ESM1b were the highest scoring unsupervised *in silico* missense predictors for IDR variants. Our baseline model, using only IDR-specific features achieved competitive performance on the hold-out test set with a PR-AUC of 0.800. Critically, when these IDR features were combined with these methods we saw significant overall improvement. The AlphaMissense-Enhanced model increased its PR-AUC from 0.807 to 0.931. Similarly, ESM1b-Enhanced improved PR-AUC from 0.679 to 0.878 and EVE increased from 0.591 to 0.918. These results demonstrate the effectiveness of our enhancements for classifying missense variants in IDRs and highlight its ability to complement existing *in silico* missense predictors.

## Introduction

Missense variants are single nucleotide polymorphisms (SNPs) that result in the substitution of a single amino acid in the protein sequence. These changes can have a wide range of effects on protein function, from benign alterations to severe pathogenic consequences. For example, the Glu6Val (E6V) variant in *HBB* is responsible for sickle cell disease[1], while the Gly380Arg (G380R) variant in *FGFR3* leads to achondroplasia[2]. Understanding the functional impact or pathogenic potential of missense variants is essential for accurate disease diagnosis, prognosis, and therapeutic decision-making. Databases such as ClinVar[3] and Human Gene mutation database (HGMD)[4] contain curated missense variants that are classified for their role in causing disease. However, a large portion of missense variants remain unclassified due to the rarity in the population and accurately predicting the deleteriousness of missense variants remains a significant challenge[5].

Many computational *in silico* missense classification predictors have been developed to aid in classifying missense variants[6-11]. These models employ machine and deep learning techniques, including supervised and unsupervised approaches, and rely on feature set based on sequence conservation, protein secondary structure and physicochemical properties to assess variant pathogenicity[12]. The American College of Medical Genetics and Genomics (ACMG) guidelines also recognize the importance of computational missense predictors, categorizing them as supportive evidence for pathogenic (PP3) and benign (BP4) classifications[13], under the assumption that a prediction of altered function is equivalent to pathogenicity[14]. Recent predictors such as AlphaMissense[15], ESM1B[16], and EVE[8] have demonstrated strong performance from missense variants in structured regions. However, the performance in intrinsically disordered regions (IDRs) are suboptimal compared to their performance in ordered regions [15, 17, 18]. For example, AlphaMissense reports an average area under the curve (AUC) of 0.94 for missense variants in ordered regions but only about 0.85 for missense variants in disordered regions.

Traditional *in silico* missense classification predictors predominantly rely on sequence conservation-based features, assessing variant tolerance in highly conserved genomic regions and their impact on secondary structure. Even though 15-20% of IDRs are in regions with low sequence conservation, they are generally less conserved than structured domains and exhibit greater tolerance to variants[19]. Unlike ordered protein domains, IDRs lack stable conformation, can adopt multiple structural states, and exist in a low-energy state, making them challenging to model using conventional computational approaches. Despite this, IDRs play essential roles in protein function, with approximately 25% of disease-associated variants localized within these regions[18].

IDRs are highly abundant in the eukaryotic proteome, accounting for nearly 30% of all proteins, and play critical roles in processes such as transcriptional regulation, DNA replication, and signal transduction[20]. IDRs are protein segments that lack a stable secondary or tertiary structure and exist as dynamic ensembles. Their structural flexibility makes them challenging to study using X-ray crystallography or cryo-electron microscopy[21]. Instead, IDRs are typically identified based on their amino acid composition, and various disorder prediction algorithms have been developed to assess their propensity for disorder[22]. AlphaFold2 demonstrated a strong correlation between low-confidence predictions and intrinsic disorder. A study found that a combination of the confidence score Predicted Local Distance Difference Test (pLDDT) from AlphaFold2 and Relative Solvent Accessibility (AlphaFold-RSA) provides a robust approach for predicting disordered regions within proteins[23, 24].

IDRs do not adopt a single stable structure; instead, they exist in multiple conformations, influencing their flexibility and interaction potential. Recent advances in the prediction of global protein conformation (gIDRc) can predict biophysical properties of IDRs[20], providing insight into their structural adaptability altered behavior. Beyond structural flexibility, IDRs often drive biomolecular phase separation (PS), forming dynamic membrane-less organelles that regulate cellular organization. Variants within IDRs can disrupt PS, leading to loss or gain of function and contributing to diseases such as neurodegeneration and cancer[18]. Several novel computational methods exist that can predict PS using different approaches. PSAP[25] uses 55 features from amino acid composition, trained on 90 manually curated human proteins. PSPHunter[26] employs 123 features including sequence embeddings and evolutionary data from large multi-species databases. catGRANULE 2.0[27] integrates 128 features combining physicochemical properties with AlphaFold2 structural information. These methods specifically predict protein condensate formation and PS propensity, that differs from traditional *in silico* missense predictors that focus on protein stability, folding, and general functional disruption. PS predictors instead assess the capacity for proteins to form dynamic, reversible liquid-like assemblies that are critical for cellular organization and regulation. In addition to biophysical modeling, protein language models (pLMs) offer a powerful approach to understanding missense variant effects in IDRs. Recent studies have demonstrated that pLMs can effectively capture sequence-based features relevant to protein function, making them valuable tools for missense variant prediction[28].

To address the challenge of missense variant classification in IDRs, we developed a machine learning model that integrates biophysical features unique to these regions. Specifically, our model incorporates predictions of gIDRc, PS, and contextual protein embeddings derived from pLMs. We first establish a baseline model using only these IDR-specific features. We then demonstrate that this framework can be used to significantly enhance the performance of existing state-of-the-art unsupervised predictors, such as AlphaMissense, EVE, and ESM1b, by integrating their scores as additional features. This work presents a novel framework that provides a more nuanced and accurate classification of missense variants in disordered regions, addressing a critical gap in variant interpretation.

## Materials and methods

### Model generation

To analyze the impact of missense variants in IDRs, we utilized protein coordinate predictions of disorder from AlphaFold-RSA, downloaded from MobiDB[29]. The dataset was filtered for human proteins (NCBI Taxon ID 9606), and reference protein FASTA sequences for all proteins were downloaded from UniProt using their corresponding UniProt IDs (Figure 1A). These reference sequences represent the full-length protein sequences, incorporating regions predicted to be disordered by AlphaFold-RSA. We then identified all missense variants from the ClinVar database that overlapped with these disordered regions. For each of these variants, a corresponding mutant FASTA sequence was generated by introducing the variant within the predicted disordered segment of the reference sequence (Figure 1B). This approach ensured that only variants occurring within IDRs were considered for further analysis. The reference and mutant FASTA sequences were then used as inputs for feature and embedding extraction using tools ALBATROSS[20], PSAP[25], and ProtTrans[28]. ALBATROSS is used to predict biophysical properties that can be used to infer global protein conformation from an IDR. PSAP is a predictor that predicts PS from an IDR protein sequence. For ALBATROSS and PSAP, the absolute delta change between the reference and mutant sequences was computed to quantify the structural and PS alterations caused by the variant. With ProtTrans, protein embeddings were initially generated for each amino acid across the entire IDR protein sequence input, subsequently, the mean of the embeddings of size 1024 across all amino acids in the sequence was calculated to create a single scalar representation for the protein sequence. This process was performed separately for the reference and mutant sequences. To capture variant induced shifts in the embeddings, the average of the reference and mutant embeddings was computed. This aggregated representation effectively summarizes the contextual changes introduced by the variant and was used as the input feature set for downstream analysis. These embeddings of size 1024 was further refined to only the top 20 embeddings that were most predictive to deleterious function. The extracted features from ALBATROSS, PSAP and ProtTrans were then concatenated. A gradient boosting classifier (XGBoost) was trained, optimized using hyperparameter tuning with Optuna[30] and validated on a hold-out test set that resulted from randomized splitting train and test to assess its performance in distinguishing the impact of missense variants in IDRs (Figure 1C). The code for this study is available at https://github.com/rohandavidg/IFP-MIDR

**Figure 1.**
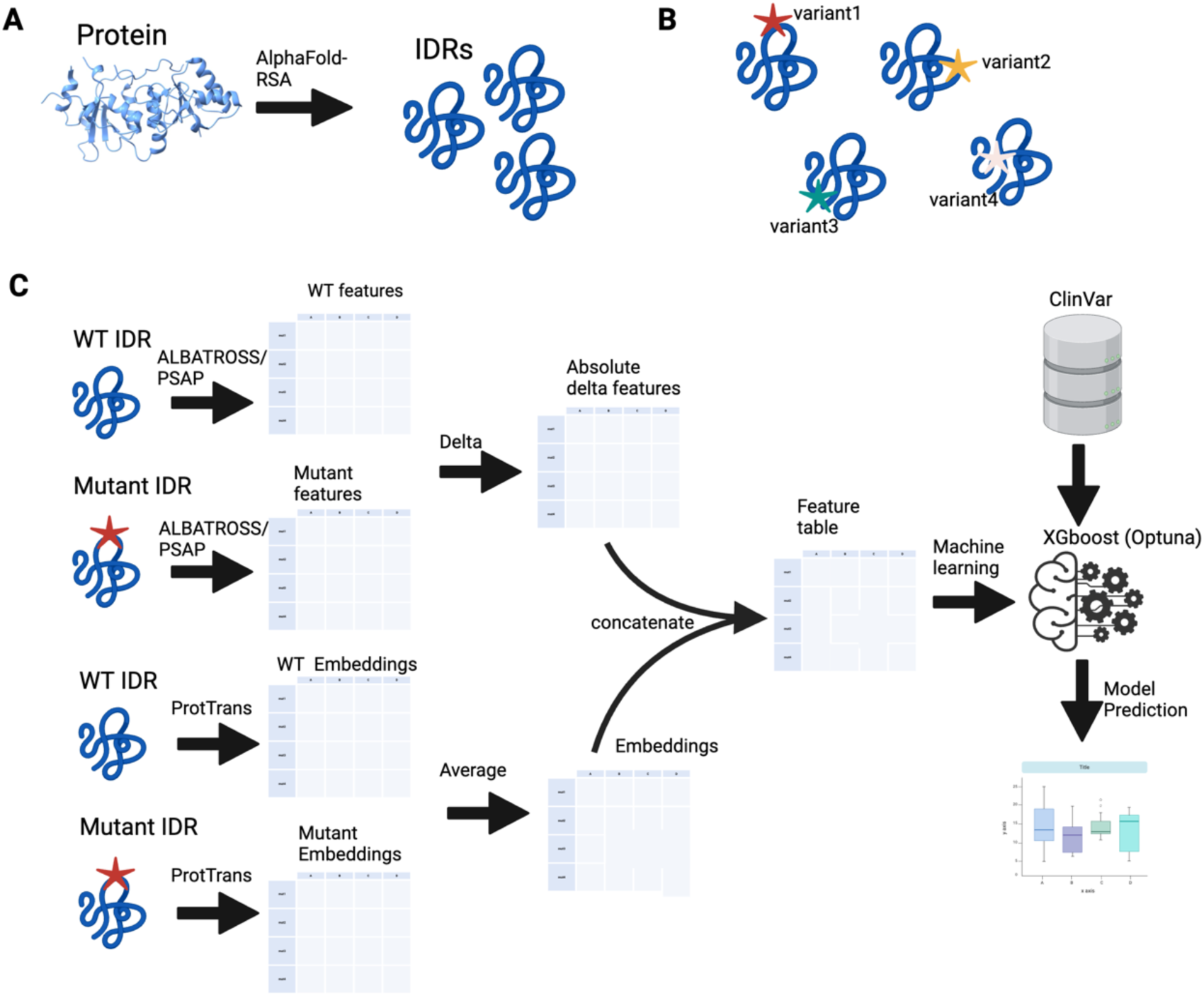
Illustration figure of the computational framework for predicting the impact of missense variants in Intrinsically Disordered Regions (IDRs). (A) Predictions from AlphaFold-RSA were used to predict the IDRs from AlphaFold structures. (B) Missense variants were introduced into the FASTA sequence of the predicted IDR sequence after mapping the protein coordinates with the genomic coordinates listed in the ClinVar database. DiJerent variants (variant1, variant2, variant3, variant4) represent variants occurring at various positions of the IDRs. (C) Features were extracted from both wild-type (WT) and mutant IDRs using ALBATROSS and PSAP to capture biophysical properties predictive of global conformation and phase separation. The absolute delta between WT and mutant features was computed. Additionally, ProtTrans embeddings were generated for both WT and mutant sequences. The embeddings were combined using the average. The extracted features and embeddings were concatenated into a feature table and used as input for an XGBoost model optimized with Optuna. The baseline model was trained using ClinVar classifications as ground truth for predictions. The final model outputs classification results evaluated using performance metrics such as AUC and PR-AUC. Created with BioRender.com.

### Data selection

The variants used in this study were downloaded from the ClinVar 2024-09-17 release, with variants located in Pfam domains filtered out as per the coordinates downloaded from University of California Santa Cruz (UCSC) resources[31]. The ClinVar VCF file was annotated with CAVA v2[32] to determine the protein substitution from genomic variants, leveraging transcripts from NCBI and EMBL-EBI (MANE) transcripts that corresponds to UniProt protein FASTA reference. IDRs predicted by AlphaFold2-RSA were observed in 834 genes that mapped to human NCBI Taxon ID 9606. To align these predictions with ClinVar transcript annotations, Ensembl transcripts associated with UniProt IDs were mapped to NCBI transcripts. This process ensured that protein coordinates were consistently aligned with genomic coordinates based on transcript information from the annotated ClinVar VCF, resulting in a total of 15,999 variants located in predicted IDR regions. Based on the clinical classification in ClinVar, 85.7% of the variants in IDRs are classified as Variant of Uncertain Significance (VUS) (Figure S1A).

The ClinVar classification of missense variants was grouped into three categories, which were Deleterious, Neutral and VUS. Variants labeled as ‘Likely_pathogenic’, ‘Pathogenic/Likely_pathogenic’, ‘Pathogenic|drug_response’, ‘Pathogenic’, ‘Pathogenic|other’, ‘Pathogenic/Likely_pathogenic|other’, ‘Likely_pathogenic|other’, and ‘Likely_pathogenic/Likely_risk_allele’ were classified as Deleterious, while those categorized as ‘Likely_benign’, ‘Benign’, and ‘Benign/Likely_benign’ were classified as Neutral, and the remaining was categorized as VUS. Further refinement to include only genes with Ensembl transcripts that could be mapped to an orthogonal RefSeq transcript reduced the dataset to a total of 2,203 variants (Figure S1B).

To evaluate the performance of existing predictors for variants found in predicted IDRs, we utilized data from dbNSFP v4.8[33], which includes a comprehensive set of *in silico* missense prediction scores from 56 predictors, including unsupervised models such as AlphaMissense, ESM1b, and EVE. Mapping all variants from the dbNSFP database to ClinVar variants located within predicted IDR regions resulted in a final dataset of 2,104 missense variants with known classifications in 290 genes (Supplementary Table S1). In total, the dataset comprised of 316 variants classified as deleterious and 1788 variants classified as neutral. The predictive performance of each *in silico* missense predictor was assessed using its normalized rank scores available in the dbNSFP database.

### Feature generation

Features such as radius of gyration, end-to-end distance, asphericity, and prefactor predicting gIDRc were generated using ALBATROSS within Sparrow v0.2.3 for both mutant and WT using the input FASTA sequences representing IDRs. Features predicting PS were evaluated using PSAP v1.0.7, which computed 15 features for wild-type (WT) and mutant sequences. To incorporate sequence level representations, we used the pLM ProtTrans, generating 1024-dimensional embeddings for both mutant and WT sequences for every amino acid. To derive a single representative embedding per sequence, the mean of the amino acid level embeddings across the entire protein chain was computed. Prior to model training, we applied supervised feature selection using F-statistic scoring to reduce protein embedding dimensions from their original high-dimensional space to 20 dimensions, retaining only the most predictive embedding features while preserving all other features and the specific standalone predictor for each model. These combined features of gIDRc, PS, and deep learning-based embeddings provided a comprehensive representation of the potential deleterious impact of missense variants in IDRs.

### Data preprocessing

The features generated by ALBATROSS and PSAP for both mutant and WT IDR sequences were used to compute the delta change, capturing the difference between the two sequence states. The absolute values of these deltas were then calculated to ignore the direction of change and quantify the magnitude of change independent of direction. Additionally, to evaluate the underlying representation of different embedding combination approaches to protein function, the pLM embeddings generated by ProtTrans for the mutant and WT sequences were calculated using four different combination strategies, that were L1 (absolute difference), L2 (Euclidean distance), average (mean of embeddings), and Hadamard product (element-wise dot product).

Let Em and Ew represent the embeddings from the mutant and WT, respectively. We define Hadamard, Average, L1 and L2 as follows:

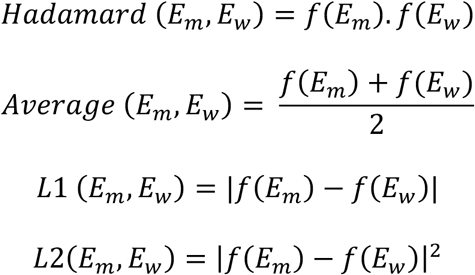

### Model creation

The features derived from ALBATROSS, PSAP, and the ProtTrans embeddings were generated for a total of 2,104 variant, comprising 316 deleterious and 1788 benign variants. These features were concatenated to form a comprehensive dataset for model evaluation. To assess predictive performance, we compared different model predictors, which were, Random Forest, Multi-Layer Perceptron (MLP), Naïve Bayes, and XGBoost classifiers using scikit-learn v0.24.2. Prior to model training, the dataset was standardized using the StandardScaler from scikit-learn, and the class labels were binarized using LabelEncoder. We employed 10-fold cross-validation to ensure robust model evaluation from an imbalanced dataset, computing key performance metrics including AUC and PR-AUC to compare model effectiveness in distinguishing deleterious from benign variants.

### Hyperparameter optimization (HPO)

We employed the Optuna framework to optimize the hyperparameters of the XGBoost model. The dataset was split into an 80:20 ratio randomly, where 80% of the data (1,683 variants) was used in 10-fold cross-validation to determine the optimal hyperparameters. In a subsequent experiment, we implemented gene-based splitting to prevent data leakage, ensuring that variants from the same gene were not present in both training and testing sets. This resulted in 1852 variants in the training and 252 variants in the test. A total of 150 optimization trials were conducted, with the objective of maximizing the PR-AUC across the 10-fold cross-validation. Once the optimal hyperparameters were identified, the final tuned model, which we term as the baseline model was applied to the hold-out test set (421 variants) to evaluate its performance on unseen data.

### Combination with AlphaMissense, ESM1b and EVE

The XGBoost model was retrained, and HPO was performed with AlphaMissense, ESM1b and EVE added separately as additional features to assess their impact on predictive performance. We term these models as AlphaMissense-enhanced (baseline and AlphaMissense), ESM1b-enhanced (baseline and ESM1b) and EVE-enhanced (baseline and EVE). The experimental design maintained consistency with our initial approach, utilizing both random and gene-based data splitting strategies. For gene-based splitting, the dataset was partitioned to prevent data leakage, ensuring variants from the same gene were not present in both training (1,852 variants) and testing (252 variants) sets. HPO employed the Optuna framework across 150 optimization trials, with the objective of maximizing PR-AUC during 10-fold cross-validation. Following optimization, final models were evaluated on the hold-out test set, with performance confidence intervals calculated through bootstrap resampling over 1,000 iterations to ensure robust statistical assessment.

### ClinVar review status classification

To assess whether prediction accuracy of the baseline model depends on variant classification quality, we stratified the hold-out test set according to ClinVar’s star-based review status system. Variants labeled as “no assertion criteria provided” or “criteria provided, single submitter” were grouped under the single-star category. Variants with “criteria provided, multiple submitters, no conflicts” were categorized as two stars, while those “reviewed by expert panel” were assigned to the three-star category. This stratification resulted in 292 variants in the one-star category, 100 in the two-star category, and 29 in the three-star category. Within each star category the PR-AUC were calculated, along with its CIs.

### Comparative analysis of dbNSFP *in silico* missense predictors and the enhanced predictors

To evaluate whether our methodology demonstrates superior performance on variants where existing missense predictors struggle, we conducted a systematic analysis of variant-level classifications across all *in silico* missense predictors in the dbNSFP database using our hold-out test set. For each IDR variant, we calculated the misclassification rate among existing predictors relative to ClinVar annotations and stratified variants by error rate to identify those that consistently challenge current computational approaches. We then assessed the performance of the baseline and enhanced models’ ability to accurately classify these difficult-to-classify variants.

### Statistical analysis

Baseline model performance was assessed using AUC and PR-AUC metrics across four machine learning algorithms: XGBoost, Random Forest, Naïve Bayes, and multilayer perceptron (MLP). The evaluation framework employed 10-fold cross-validation for model training and hyperparameter optimization, followed by performance assessment on an independent hold-out test set to ensure unbiased evaluation.

The association between individual biophysical features and variant pathogenicity was assessed using the non-parametric Mann-Whitney U test. To evaluate statistical significance of performance differences between models, we compared the distributions of AUC and PR-AUC scores across cross-validation folds using the Mann-Whitney U test. This approach enabled pairwise comparisons between the baseline model, enhanced models and standalone *in silico* predictors (AlphaMissense, ESM1b, EVE), and final enhanced models. For all performance metrics on the independent test set, 95% confidence intervals were calculated using bootstrap resampling with 1,000 iterations.

## Results

### Evaluation of dbNSFP predictors with functions

A significant number of *in silico* missense predictors are trained on ClinVar and HGMD datasets, which exhibit substantial overlap in their training data and the data used in this study. However, 23 models in the dbNSFP database are not trained on ClinVar and HGMD. Among these, when focusing on variants located in predicted IDR regions, AlphaMissense, an unsupervised model, demonstrated the highest predictive performance with an AUC of 0.908 (95% CI:0.889-0.924) (Figure 2A) and PR-AUC of 0.733 (95% CI:0.690-775) (Figure 2B), followed by ESM1b with an AUC of 0.847 (95% CI: 0.819–0.872) and PR-AUC of 0.606 (95% CI:0.548-0.662). Additionally, EVE, the other unsupervised autoencoder model using sequence conservation, achieved an AUC of 0.715 (95% CI: 0.679–0.751) and PR-AUC of 0.50 (95% CI:0.446-0.556), highlighting its moderate performance for missense classification in predicted IDRs. These results, derived from ClinVar-labeled variants located in AlphaFold-RSA predicted disordered regions, suggest that AlphaMissense currently represents the state-of-the-art benchmark for predicting deleterious missense variants in IDRs (Supplementary Table S2).

**Figure 2:**
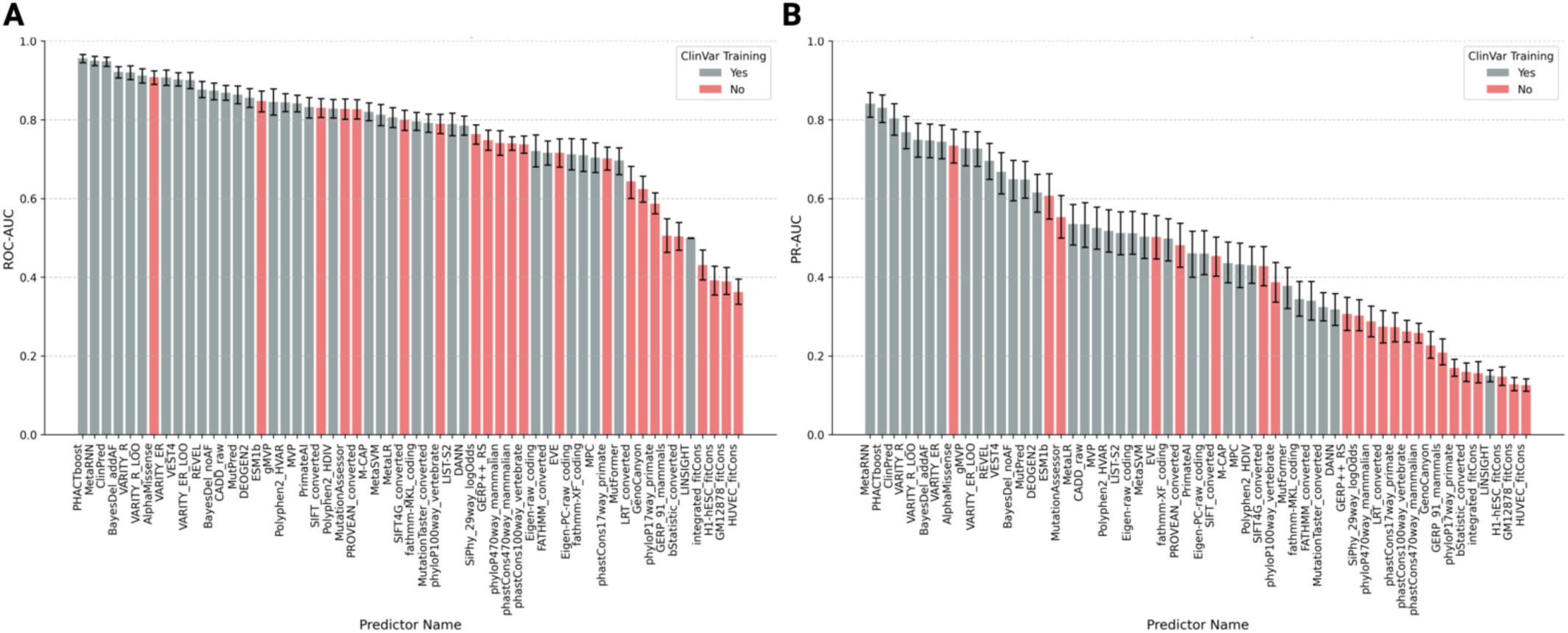
Performance metrics of In Silico missense predictors on variants found in IDR regions A: The area under the curve performance of all in silico missense predictors listed in the dbNSFP databases for variants found in the ClinVar database. The predictors in red highlight those that have not been trained using ClinVar labels. B: Precision-Recall performance for the same subsets.

### Association of global IDR conformation with protein function

Using a Mann-Whitney U test, we evaluated the association between protein function and the absolute changes in radius of gyration, end-to-end distance, asphericity, and prefactor between the WT and mutant sequences. All four features demonstrated a significant association with protein function. To examine their distribution across deleterious and neutral categories, a square root transformation was applied to all four features (Figure 3A). Further evaluation of the predictive performance based on the AUC, revealed that the absolute change in scaling exponent had the highest AUC of 0.628. Notably, all five features achieved an AUC greater than random prediction (Figure 3B), highlighting their potential relevance in impact assessment.

**Figure 3:**
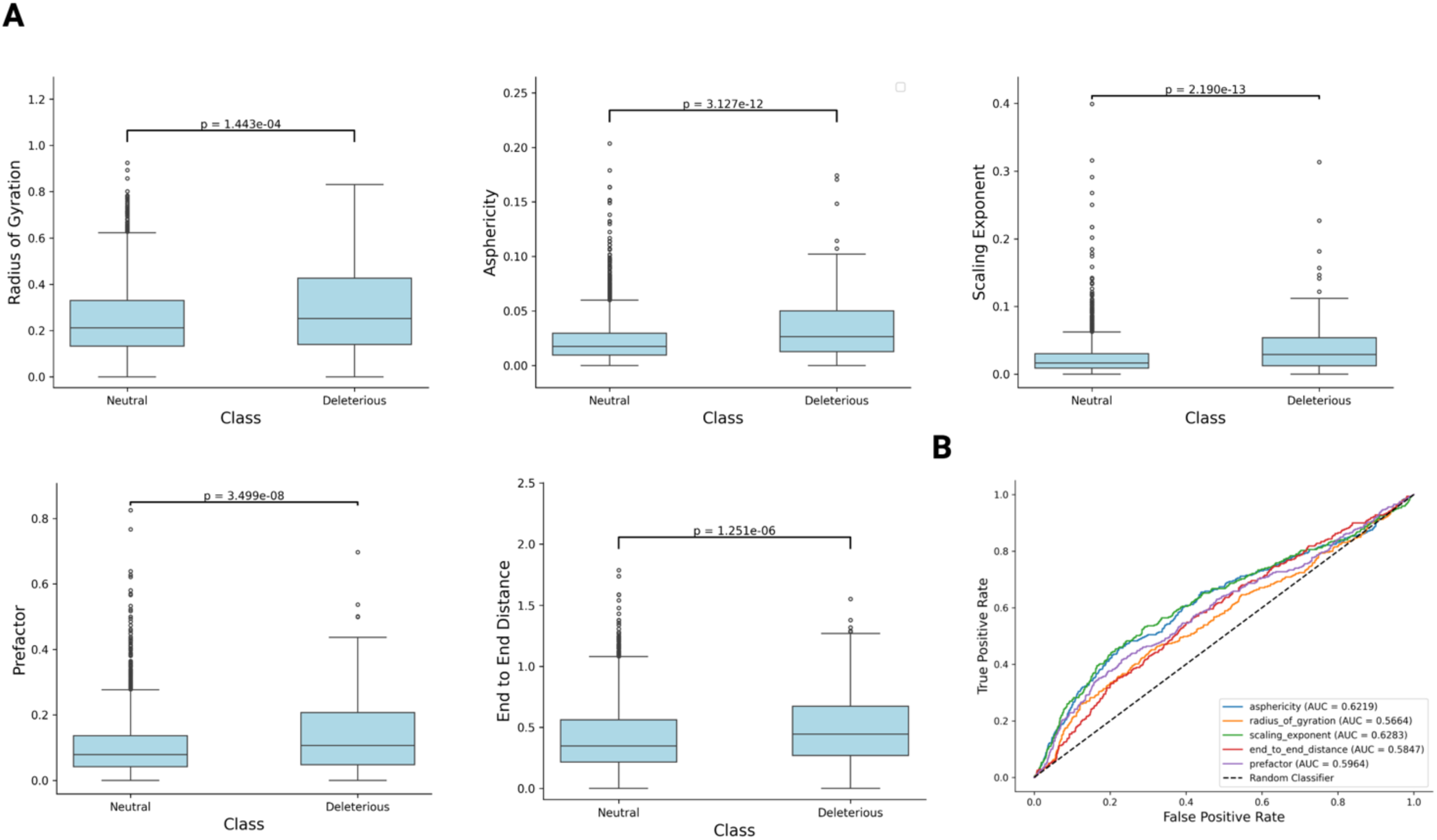
Absolute change in global protein conformation due to a missense variant in an IDR contributes significantly to protein prediction. (A) Association of the absolute change in Radius of Gyration, Asphericity, Scaling Exponent, Prefactor, End to End distance transformed using the square root between the reference and mutant with protein function. Outliers were removed to observe the overall distribution. (B) Area under the curve performance highlighting the prediction of protein function from absolute change in Radius of Gyration, Asphericity, Scaling Exponent, Prefactor, End to End distance induced due to a missense variant.

### Evaluation of embedding combination method and model performance

We evaluated four different embedding combination methods (L1, L2, average, and Hadamard) across four different machine learning and deep learning models (XGBoost, Random Forest, Naïve Bayes, and MLP) to predict protein function Using PR-AUC and AUC across 10-fold cross-validation, we found that the average and Hadamard combination methods, when integrated with features from PS and gIDRc and modeled using either XGBoost or Random Forest, achieved the highest predictive performance. Specifically, with XGBoost, both the average and Hadamard methods yielded high mean AUC values of 0.904 and 0.908, respectively, with no statistically significant difference between them (Figure 4A and Supplementary Table S3). Similarly, both methods achieved a mean PR-AUC of 0.76 and 0.74 with XGBoost, again showing no significant difference (Figure 4B). When evaluated using Random Forest, the average and Hadamard methods also exhibited comparable performance. However, models utilizing Hadamard and average embeddings with XGBoost and Random Forest significantly outperformed those using L1 and L2 embeddings, demonstrating statistically significant improvements in both AUC and PR-AUC. The default hyperparameters for XGBoost included learning_rate=0.3, n_estimators=100, max_depth=6, gamma=0, and subsample=1.0. Given its robust performance, we selected XGBoost trained on the average embedding combination, along with features from gIDRc and PS, as the final model for further optimization.

**Figure 4:**
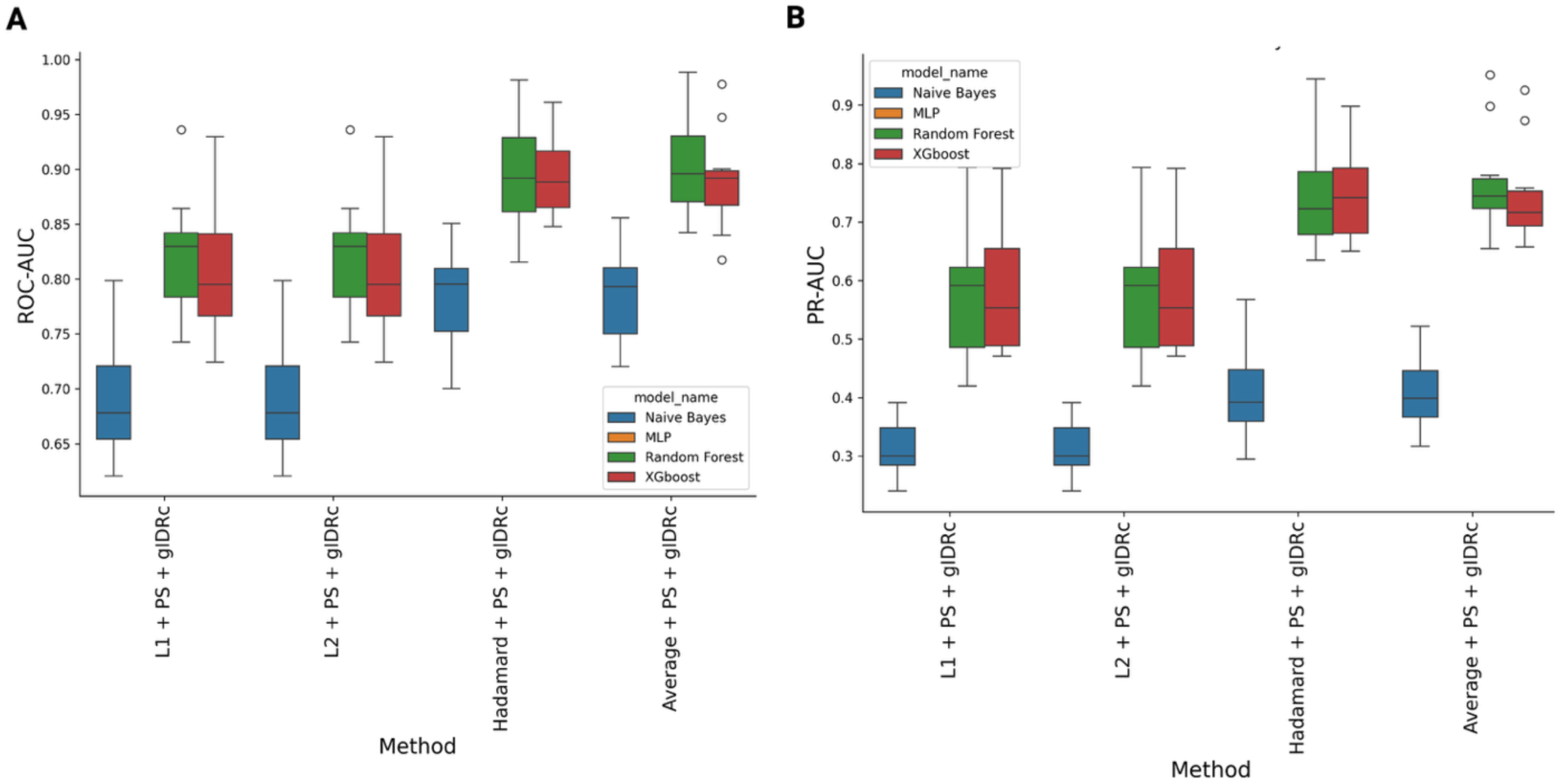
Performance comparison of multiple protein embedding combination methods using multiple machine learning models to predict protein function on classified variants found in ClinVar in predicted IDR regions. (A) Area under the curve performance of XGBoost, Random Forest, Naïve Bayes and Multi-Layer Perceptron using protein language model embeddings (L1, L2, average and Hadamard), along with features predicting absolute change in Phase Separation and absolute change in global IDR conformation to predict protein function from a missense variant. (B) Area under the precision-recall curve performance of XGBoost, Random Forest, Naïve Bayes and Multi-Layer Perceptron using protein language model features (L1, L2, average and Hadamard), along with features predicting absolute change in Phase Separation and absolute change in global IDR conformation to predict protein function from a missense variant.

### Hyperparameter optimization (HPO)

Using the Optuna framework, we further optimized the XGBoost model through 150 trials, aiming to maximize the mean AUC across 10-fold cross-validation. Each trial represents a new set of hyperparameters that is tested. HPO resulted in an increase in mean PR-AUC from 0.802 to 0.840, demonstrating a slight but meaningful improvement in predictive performance from the training set. When applied to the hold-out test set, which comprised of 421 unique variants, the optimized model achieved an PR-AUC of 0.800 (95% CI 0.7113-0.876), which we refer to as the baseline model. To ensure there was no data leakage, we computed pairwise sequence identity between sequences in the training and test sets and found that 0% of sequences in the test set had 100% identity with those in the training set. The final set of optimal hyperparameters identified included: n_estimators=330, max_depth=10, learning_rate=0.077, colsample_bytree=0.772, subsample=0.505, gamma=624, min_child_weight=3, reg_alpha=1.68, and reg_lambda=5.14. Among these, “gamma”, “reg_alpha”, and “min child weight” were found to be the most influential and showed the strongest correlations with PR-AUC performance. The negative correlations indicate that lower regularization and reduced constraints on tree splitting improved model performance, suggesting the model benefits from increased flexibility rather than regularization constraints. (Figure S2).

### Feature evaluation

Utilizing SHapley Additive exPlanations (SHAP)[34] to assess feature importance in the optimized XGBoost model revealed that the embeddings combined via the average method between the WT and mutant were the most influential (Figure S3). The analysis demonstrated that pLM embeddings from ProtTrans constituted the most influential features, with the highest-ranking feature achieving a SHAP importance value of approximately 0.09. PS features from PSAP and gIDRc features from ALBATROSS were prominently distributed throughout the importance hierarchy, with several features from each category ranking within the top 20 most influential predictors. The balanced representation of features from ProtTrans, PSAP, and ALBATROSS within the top performing features validates our integrative approach, demonstrating that pLM embeddings, PS propensity, and gIDRc provide distinct yet synergistic information for predicting missense variant pathogenicity.

### Improvement with AlphaMissense, ESM1b and EVE

As noted above, AlphaMissense, ESM1b and EVE were the highest performing unsupervised classification models. To improve these metrics, we retrained the XGBoost model and optimized its hyperparameters to create an “Enhanced” version. Enhanced model creation begins with one of AlphaMissense, ESM1b or EVE. Next, PS, gIDRc and the average of the embeddings between the WT and mutant are added. The dataset was split into 80% training data (1,683 variants), where 10-fold cross validation was performed, and 20% (421 variants) was reserved as a hold-out test set.

EVE incorporation as a feature also demonstrated significant performance gains. The EVE-enhanced model achieved a mean PR-AUC of 0.813 in 10-fold cross-validation, substantially outperforming standalone EVE (PR-AUC: 0.506; 95% CI: 0.475-0.537; p < 0.001; Cohen’s d = 5.46). Hold-out test validation showed EVE-enhanced achieving an PR-AUC of 0.918 (95% CI: 0.860-0.962) compared to standalone EVE, with a PR-AUC of 0.591 (95% CI: 0.471-0.705; p < 0.001; Cohen’s d = 6.774) (Figure 5A). Optimal hyperparameters were n_estimators=466, max_depth=10, learning_rate=0.029, colsample_bytree=0.863, subsample=0.602, gamma=0.877, min_child_weight=1, reg_alpha=0.38, and reg_lambda=7.66.

**Figure 5:**
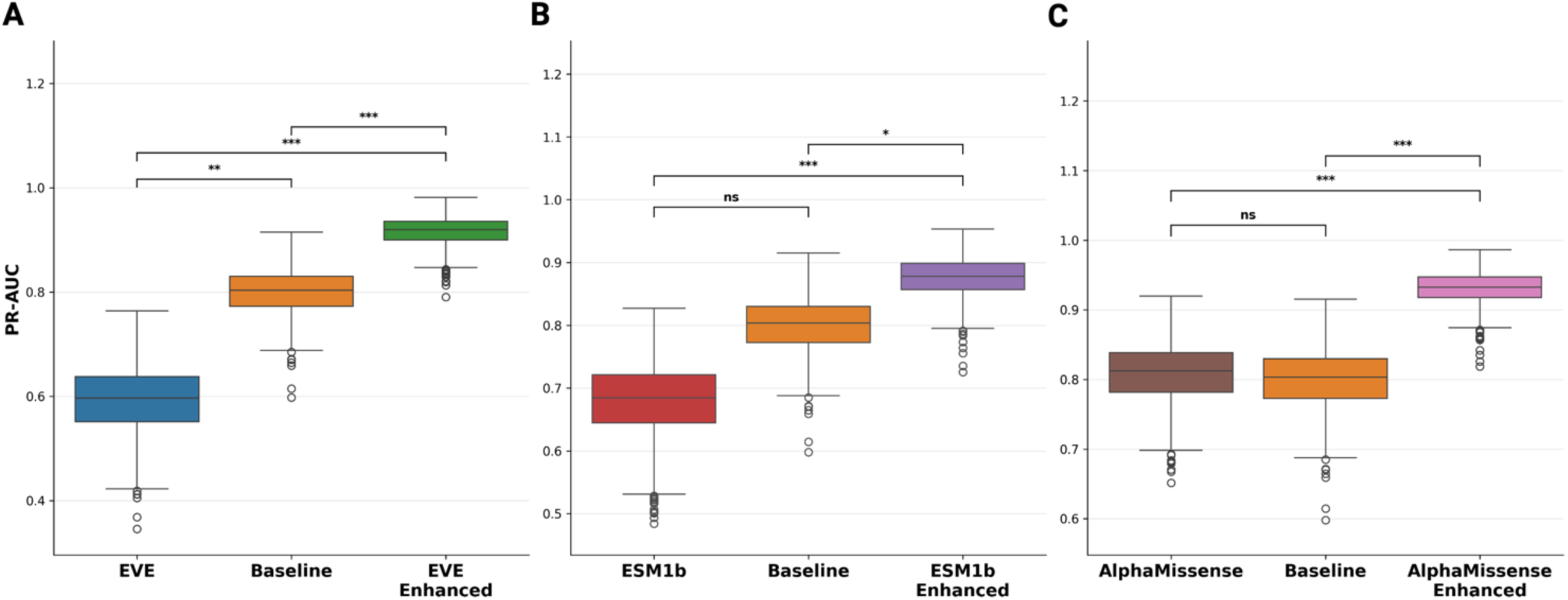
Performance comparison of protein variant eFect prediction methods. Precision-Recall Area Under the Curve (PR-AUC) metrics comparing standalone predictors versus baseline and enhanced models across EVE, ESM1b, and AlphaMissense methods on the hold-out test set. **(A)** PR-AUC performance with confidence intervals using bootstrap resampling with 1,000 iterations comparing EVE vs Baseline vs EVE enhanced. (B) PR-AUC performance with confidence intervals using bootstrap resampling with 1,000 iterations comparing ESM1b vs Baseline vs ESM1b enhanced (C) PR-AUC performance with confidence intervals using bootstrap resampling with 1,000 iterations comparing AlphaMissense vs Baseline vs AlphaMissense enhanced. Statistical significance was assessed using pairwise Mann-Whitney U tests: ***p < 0.001, **p < 0.01, *p < 0.05, ns = not significant.

ESM1b integration demonstrated equally substantial performance enhancement. The ESM1b-enhanced model achieved a mean PR-AUC of 0.840 in 10-fold cross-validation from the training set, significantly outperforming standalone ESM1b (PR-AUC: 0.622; 95% CI: 0.561-0.682; p < 0.001; Cohen’s d = 3.25). The 35.19% increase in PR-AUC substantially enhances deleterious variant detection capability. Hold-out test validation confirmed superior performance with PR-AUC of 0.877 (95% CI: 0.809-0.929) versus standalone ESM1b PR-AUC of 0.679 (95% CI: 0.563-0.780; p < 0.001; Cohen’s d = 4.35) (Figure 5B). Optimal hyperparameters included: n_estimators=330, max_depth=10, learning_rate=0.077, colsample_bytree=0.772, subsample=0.505, gamma=0.624, min_child_weight=3, reg_alpha=1.68, and reg_lambda=5.14.

The AlphaMissense-enhanced model achieved a mean PR-AUC of 0.863 representing a significant improvement over standalone AlphaMissense (PR-AUC: 0.740, 95% CI: 0.680-0.797; p = 0.0017, Cohen’s d = 1.84) from the training set. The 16.78% improvement in PR-AUC is clinically significant as it directly measures enhanced detection of deleterious variants. Independent validation on the hold-out test set confirmed this enhancement, with PR-AUC improving from 0.807 (95% CI: 0.715-0.885) to 0.931 (95% CI: 0.881-0.971; p < 0.001; Cohen’s d = 3.882), representing a 16.52% improvement (Figure 5C). This performance rivals top *in silico* missense predictors trained on ClinVar variants, including MetaRNN (PR-AUC: 0.841, 95% CI: 0.806-0.869) and PHACTboost (PR-AUC: 0.830, 95% CI: 0.793-0.863). Optimal hyperparameters were determined through Optuna optimization: n_estimators=390, max_depth=3, learning_rate=0.068, colsample_bytree=0.83, subsample=0.595, gamma=1.50, min_child_weight=1, reg_alpha=1.12, and reg_lambda=1.192.

### Model Performance on ClinVar variants based on review status

Overall, the baseline model achieved a PR-AUC of 0.800 (95% CI 0.711-0.876) on a hold-out test set comprising 421 variants. Stratification by ClinVar review status revealed quality-dependent performance patterns, with one-star variants (lowest evidence quality, n=292) achieved PR-AUC of 0.724 (95% CI: 0.592-0.833), two-star variants (moderate evidence quality, n=100) achieved higher PR-AUC of 0.824 (95% CI: 0.639-0.945), and three-star variants (expert panel review, n=29) achieved PR-AUC of 0.700 (95% CI: 0.100-1.000). The performance estimate for three-star variants has substantial uncertainty, as evidenced by the confidence interval spanning nearly the entire possible range due to the limited sample size. Better performance on two-star variants suggests the model performance improved on variants with better evidence quality (Figure S4).

### Comparative Analysis of *In Silico* Missense Predictors and Enhanced Model Performance

Evaluation of our baseline model against established *in silico* predictors revealed superior performance on challenging variants within the hold-out test dataset. Several variants showed consistently high error rates across most predictors, highlighting the challenge in predicting these variants with high accuracy. Baseline models correctly classified several variants that were consistently misclassified by the majority of dbNSFP predictors. For instance, likely pathogenic variants R573C in FGA and A654V in HIF1A were incorrectly predicted as neutral by most existing tools, including AlphaMissense, while our baseline model accurately identified them as deleterious. Conversely, likely benign variants T319M and P869S in GLI3, and S186Y in BRCA1, were incorrectly predicted as damaging by many *in silico* predictors, whereas the baseline model correctly classified them as neutral. These results demonstrate the baseline models’ capability to handle challenging variants that represent common failure modes for existing prediction tools (Supplementary Table S4).

Enhanced models demonstrated substantial improvements over their standalone counterparts across all three integrated predictors. AlphaMissense-Enhanced correctly classified 32 variants that were misclassified by standalone AlphaMissense, including 8 pathogenic/likely pathogenic and 24 benign/likely benign variants. Similarly, ESM1b-Enhanced accurately classified 28 variants missed by standalone ESM1b, comprising 10 pathogenic/likely pathogenic and 18 benign/likely benign variants. EVE-Enhanced model showed the most substantial improvement, correctly identifying 47 variants that standalone EVE misclassified (27 pathogenic/likely pathogenic and 20 benign/likely benign variants) using an optimal cut-point of 0.028. These results demonstrate consistent enhancement across all integrated predictors, with strength in reducing false negative classifications of pathogenic variants while maintaining specificity for benign variants.

## Discussion

The relatively lower predictive performance of *in silico* missense classification in IDRs, compared to structured domains is well-documented[15, 17]. This discrepancy may stem from the fact that most known pathogenic missense variants reside in ordered protein regions. Due to their lack of well-defined secondary structures, low evolutionary conservation, and highly dynamic nature, variants in IDRs present significant challenges for accurate prediction of missense classification. Many computational models are overfit to structured regions, leading to biased predictions that underperform for disordered region variants. Given these challenges, new computational strategies incorporating biophysical properties unique to disordered regions are necessary to enhance the classification of missense variants within IDRs.

PLMs like ProtTrans, combined with biophysical predictors such as Albatross and PSAP, offer a powerful approach to capturing biophysical information embedded within protein sequences, providing valuable insights into predicted consequences. The optimal method for integrating embeddings from WT and mutant sequences remains system dependent. In this study, we found that the Hadamard product and average embedding combination methods outperformed L1 and L2 distance-based approaches, demonstrating their effectiveness in assessing the impact of missense variants in IDRs.

Recent studies have shown an association between changes in PS and overall protein function[18]. Our findings further highlight that alterations in gIDRc also contribute significantly to predictions, though it is not the sole predictor. Notably, we demonstrate that gIDRc, PS, and pLM embeddings create a robust framework for predicting the consequences of missense variants in IDRs. This approach significantly outperforms leading *in silico* missense models, including ESM1b and EVE.

Moreover, incorporating AlphaMissense, ESM1b, or EVE as additional features further enhances predictive accuracy, suggesting that gIDRc, PS, and protein embeddings provide independent and complementary information. Feature importance analysis using SHAP further supports this conclusion, revealing that gIDRc, PS, and protein embeddings rank among the top predictive features, underscoring their critical role in classification of IDR associated missense variants. The improvement in prediction classification performance is now comparable to top *in silico* missense classification methods trained directly on ClinVar annotations.

Our study has several limitations that provide avenues for future work. First, our dataset relied on computational predictions of disorder by AlphaFold-RSA rather than experimentally validated IDRs. Although filtering variants within known Pfam domains increased confidence in our disorder predictions, this approach reduced dataset size and precluded comprehensive validation against experimentally confirmed IDRs. Notably, only 18 neutral variants in our hold-out test set were confirmed to reside within validated DisProt regions, all of which were correctly predicted by the three enhanced models. The use our gene-based data splitting strategy, while necessary to prevent information leakage, resulted in a relatively small and heterogeneously distributed test set that limited the statistical power of our evaluation. Future studies would benefit from larger, more balanced datasets that incorporate experimentally validated disordered regions.

Second, the general sparsity of high-confidence, experimentally annotated pathogenic variants in IDRs presents a major challenge for training and robustly evaluating any predictive model. To address this, future studies could employ semi-supervised learning to leverage the vast amount of unlabeled variant data or use generative models to augment training sets with biologically plausible synthetic variants.

Finally, there is a remaining knowledge gap in the specific molecular mechanisms by which IDR variants exert their effects, such as altering post-translational modifications, cellular signaling, or protein self-assembly. Further investigation into these downstream consequences is necessary to fully understand the functional impact of variation in disordered regions.

While computational predictors have significantly improved missense variant classification, major limitations persist in their ability to assess IDRs. Traditional models overemphasize evolutionary conservation and structural stability, resulting in biased predictions that fail to capture the complexity of IDRs. Our baseline model introduces an IDR specific framework that integrates predictions of global conformation changes, PS dynamics, and deep learning-based embeddings to refine missense variant classification in IDRs. By addressing the shortcomings of existing approaches, this study advances the accurate classification of IDR variants and enhances our understanding of their role in disease.

## Ethics approval and consent to participate

Not applicable

## Data Availability

The relevant code resource and datasets can be downloaded from GitHub (https://github.com/rohandavidg/IFP-MIDR). The dataset were derived from sources in the public domain: ClinVar (https://ftp.ncbi.nlm.nih.gov/pub/clinvar/vcf_GRCh37/), dbNSFP(https://www.dbnsfp.org/download), MobiDB(https://mobidb.org/)

## Author’s information

Rohan Gnanaolivu

Department of Quantitative Health Sciences, Mayo Clinic, Rochester, MN, USA

Steven Hart

Department of Quantitative Health Sciences, Mayo Clinic, Rochester, MN, USA & Department of Laboratory Medicine and Pathology, Mayo Clinic, Rochester, MN, USA

